# Reinforcement Learning for Efficient Locomotion of Bio-Inspired Microswimmers in Low Reynolds Number Fluids

**DOI:** 10.1101/2025.07.14.664791

**Authors:** Hesam Mozafarinia, Ali Zabetnia

## Abstract

Microswimmers, microscopic self-propelled agents, play vital roles in biological processes such as cellular motility, chemotaxis, and biofilm formation, and inspire the design of artificial microrobots for targeted drug delivery, microsurgery, and environmental sensing. Operating at low Reynolds numbers, where viscous forces overwhelmingly dominate and inertial effects are negligible, microswimmers face unique hydrodynamic constraints. According to Purcell’s Scallop Theorem, time-reversible or reciprocal motions cannot produce net locomotion, necessitating complex, non-reciprocal deformation cycles for effective propulsion.

In this study, we couple low Reynolds number hydrodynamics, modeled via the Stokes equation and Oseen tensor, with reinforcement learning (RL) algorithms to autonomously discover optimal swimming gaits across a range of biologically inspired microswimmer models. Beginning with the classical Najafi-Golestanian three-sphere swimmer, whose cyclical deformation mimics flagellar or ciliary beating, we validate our framework by reproducing established propulsion patterns. We then enhance biological realism by replacing rigid linkers with oscillatory elastic springs and incorporating electrostatic interactions among charged spheres, modeling chemotactic responses and intercellular electrostatic effects observed in microorganisms.

Extending beyond linear architectures, we investigate more complex geometries inspired by microbial shapes and locomotion mechanisms, including triangular, plus-like, and cage-like swimmers that emulate bacterial flagella bundling, amoeboid deformation, and cargo encapsulation strategies. Our reinforcement learning approach reveals emergent locomotion strategies reminiscent of natural traveling-wave patterns, phase synchronization, and directional taxis. Remarkably, the cage-like swimmer exhibits nearly triple the displacement of the classical three-sphere model, emphasizing the influence of geometric constraints and structural complexity on swimming efficiency.

This interdisciplinary work bridges biophysical modeling and machine learning, providing insights into microscale motility mechanisms and advancing the engineering of biomimetic microrobots capable of navigating complex fluidic environments in biomedical and ecological contexts.

## 1 Introduction

The ability to self-propel in viscous fluid environments is a fundamental requirement for many microorganisms and plays an increasingly important role in the development of artificial microscale systems. Microswimmers, whether biological or synthetic, operate in fluid regimes where the Reynolds number is extremely low, resulting in a dominance of viscous forces and a near-complete absence of inertia. This imposes strict physical constraints on propulsion mechanisms that differ significantly from those at macroscopic scales. In particular, motions that are symmetric and time-reversible fail to generate net displacement, as dictated by Purcell’s Scallop Theorem [21]. To overcome these limitations, many biological microswimmers rely on periodic, non-reciprocal deformations of their shape or internal structure to achieve locomotion. Examples include flagellar undulations in sperm cells [12], ciliary beating in protozoa [3], and traveling wave deformations observed in filamentous cyanobacteria [9]. These mechanisms provide elegant solutions evolved under the constraints of low Reynolds number hydrodynamics, offering inspiration for both theoretical modeling and the design of artificial microrobots [7].

Over the past two decades, simplified theoretical models have provided valuable insights into the basic principles of microscale propulsion. Among them, the Najafi-Golestanian three-sphere swimmer has emerged as a widely studied minimal model that captures the essential ingredients of low Reynolds number swimming through hydrodynamic interactions [19]. This model, consisting of three spherical bodies connected by extensible linkers, demonstrates how cyclic, non-reciprocal deformations can translate into net propulsion in a viscous fluid.

While these theoretical models provide critical insights, the exploration of novel swimmer architectures and their optimal control strategies remains an open problem, particularly when considering geometries beyond simple linear chains [10]. Analytical approaches become increasingly intractable as swimmer complexity grows, motivating the integration of data-driven methods with hydrodynamic modeling. Recent advances in machine learning, particularly in reinforcement learning (RL), offer promising tools to address these challenges [25]. RL has been successfully applied to a variety of physical systems to enable agents to learn optimal control policies through interaction with their environment, without explicit programming of the desired behaviora. When combined with physically accurate models of low Reynolds number hydrodynamics, RL provides a powerful framework for exploring swimmer design and control in a systematic, autonomous manner [15, 22].

In this work, we address this need by integrating low Reynolds number hydrodynamics with reinforcement learning (RL) to explore and optimize the locomotion of diverse microswimmers. Using the Stokes equation and the Oseen tensor to model hydrodynamic interactions [16], we investigate swimmer designs ranging from classical linear chains to new architectures, including triangular, plus-like, and cage-like structures. To validate our approach, we first reproduce well-established results from the literature. Specifically, our RL framework successfully recovers the traveling wave-like swimming strategies observed in filamentous Cyanobacteria for linear swimmers [9], consistent with previous theoretical predictions [13]. Additionally, we replaced the rigid linkers with oscillating springs [6] and later introduced electrostatic interactions by charging the spheres, approximating more realistic chemotactic microrobots. Previous theoretical studies have indicated that optimal propulsion in oscillatory microswimmer models typically occurs when the phase difference between linker deformations approaches *π/*2 [9, 13]. In our reinforcement learning experiments, we observed that as the swimmer models were progressively modified to more closely resemble real systems, such as by introducing elastic and electrostatic interactions, the learned phase difference converged increasingly toward this theoretically optimal value.

Building upon these validated models, we extend our investigation to more complex geometries. Our RL framework autonomously identifies efficient swimming patterns for triangular [8], plus-like, and cage-like swimmers, each exhibiting distinct locomotion capabilities. In particular, we find that the triangular swimmer moves along hexagonal trajectories with rotational flexibility, while the cage-like swimmer, constrained to linear motion, demonstrates robust learning performance. The plus-like swimmer, due to its increased geometric complexity, requires longer training but ultimately achieves directed locomotion toward target regions. Our results demonstrate the potential of combining hydrodynamic modeling with machine learning to autonomously discover efficient propulsion strategies for both classical and novel microswimmer designs. This approach bridges theoretical modeling, biological inspiration, and machine learning, contributing to the advancement of autonomous microrobotics in low Reynolds number environments [20].

## 2 Methodology

The geometric framework considered in this study is based on an extensible chain of spherical components connected by deformable linkers. Specifically, the swimmer consists of *N* rigid spheres of radius *R*, interconnected by *N* − 1 adjustable-length rods. The length of each linker, denoted by *l*, can be actively modulated, representing internal shape changes essential for propulsion. By restricting each rod to discrete states corresponding to its minimum and maximum allowable lengths, the swimmer possesses 2^*N*−1^ distinct structural configurations. Transitions between these states occur via controlled deformation of individual linkers, with each state allowing *N* − 1 possible single-linker transitions.

In the following sections, we detail the governing hydrodynamic framework describing interactions with the surrounding viscous fluid, present various swimmer architectures explored in this work, and introduce the reinforcement learning framework employed to enable autonomous discovery of efficient propulsion strategies.

### 2.1 Governing Hydrodynamics

The interaction between the microswimmer and the surrounding fluid is governed by Stokes flow, which describes viscous-dominated motion at microscopic scales. Starting from the Navier-Stokes equation [14, 27],

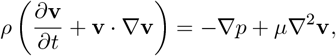

where *ρ* is the fluid density, **v** is the velocity field, *p* is the pressure, and *µ* is the dynamic viscosity.

At low Reynolds numbers, typical of microscale systems, inertial terms are negligible compared to viscous forces. Specifically, for Re ≪ 1, we have,

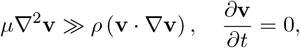

which simplifies to the steady-state Stokes equation,

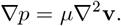

For incompressible flows, the continuity condition applies,

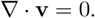

Microswimmers consists of *N* rigid spheres, each with radius *R*, connected by deformable linkers. Hydrodynamic interactions between spheres are modeled using the Oseen tensor approximation, valid for dilute systems where *R/𝓁*≪ 1, with *𝓁* the typical separation distance between spheres.

The linearity of Stokes flow allows the velocities of the spheres to be expressed as:

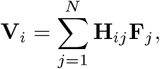

where **V**_*i*_ and **F**_*j*_ are the velocity and force acting on spheres *i* and *j*, respectively. The Oseen tensor **H**_*ij*_ is defined as:

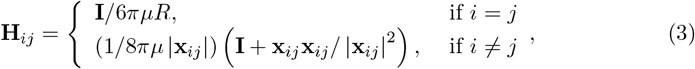

where **I** is the identity matrix and **x**_*ij*_ = **x**_*j*_ − **x**_*i*_ is the vector connecting spheres *I* and *j*.

Self-propelled motion is modeled by enforcing force-free and torque-free conditions:

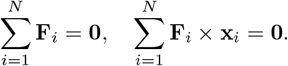

Due to the linearity and time independence of Stokes flow, net swimmer displacement depends solely on the sequence of internal shape changes, independent of deformation speed. Thus, the swimmer’s propulsion strategy is determined by the ordered sequence of configuration changes.

### 2.2 System Parameters

All swimmer models use identical physical parameters for consistency. The spheres have radius *R* = 1 *µ*m, linkers extend between *l*_min_ = 10 *µ*m and *l*_max_ = 20 *µ*m, and linker deformation speed is *u* = 10 *µ*m*/*s. The surrounding fluid has viscosity *µ* = 10^−3^ Pa · s, representing a highly viscous environment typical of microswimmer applications.

### 2.3 Swimmer Models and Architectures

The microswimmers considered in this study consist of *N* spherical bodies of radius *R*, connected by *N* − 1 extensible linkers that actively control the swimmer’s geometry. The length of each linker, *l*_*i*_, can alternate between two discrete states: a contracted length *l*_min_ and an extended length *l*_max_. This simplified binary deformation scheme provides 2^*N*−1^ possible configurations for each swimmer, with transitions corresponding to shape changes driving propulsion.

We explore both previously established and novel swimmer architectures1:

- **Linear three-sphere swimmer(Najafi-Golestanian model)**: The classical minimal swimmer model consisting of three aligned spheres and two extensible linkers, widely studied in low Reynolds number locomotion.
- **Double spring model**: An extension of the three-sphere swimmer where rigid linkers are replaced by elastic springs capable of controlled, oscillatory deformations, enabling investigation of continuous propulsion cycles.
- **Charged spring model**: A further refinement of the Double spring model in which the left sphere carries charge −2*Q*, the central sphere +*Q*, and the right sphere −*Q*, introducing electrostatic repulsion that mimics interactions present in biological or chemotactic microswimmers.
- **Generalized linear** *N* **-sphere swimmer**: A generalization of the Najafi-Golestanian model with *N* aligned spheres and *N* − 1 linkers, allowing more complex deformation cycles.
- **Triangular swimmer**: A closed-loop geometry with three spheres connected to form an equilateral triangle using three extensible linkers, enabling planar, rotational, and translational motion.
- **Plus-like swimmer**: A novel design with five spheres arranged in a cross or “plus” configuration, connected by four orthogonal linkers, introducing geometric complexity and multi-directional capabilities.
- **Cage-like swimmer**: A three-dimensional structure with a triangular cross-section, composed of multiple spheres forming a closed cage, restricted to controlled linear locomotion.

Each swimmer performs cyclic transitions between discrete configurations, with their propulsion determined entirely by the sequence of these shape changes. This framework allows systematic exploration of locomotion strategies across different geometries using reinforcement learning.

### 2.4 Reinforcement Learning Framework

To enable microswimmers to autonomously discover efficient propulsion strategies, we implement a reinforcement learning (RL) approach based on the Q-learning algorithm. In this framework, the swimmer interacts with its environment by cyclically performing internal deformations and observing the resulting displacement, progressively improving its decision-making through trial and error.

Q-learning is a model-free, off-policy algorithm designed for problems with discrete states and actions [28]. The swimmer’s possible configurations correspond to discrete states *s*, while actions *a* represent transitions between these configurations, typically by extending or contracting individual linkers. The algorithm maintains a Q-matrix *Q*(*s, a*) 2, where each entry estimates the expected cumulative reward for taking action *a* in state *s*. After each learning step *n*, the Q-matrix is updated according to [26],

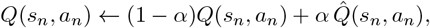

With

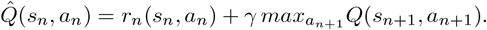

Here:

- *s*_*n*_ and *s*_*n*+1_ are the swimmer’s current and next discrete states,
- *a*_*n*_ is the chosen action at step *n*,
- *r*_*n*_ is the immediate reward based on physical displacement,
- *α* (0≤ *α*≤ 1) is the learning rate controlling how new information updates the Q-matrix,
- *γ* (0 ≤ *γ* ≤ 1) is the discount factor balancing immediate versus future rewards.

The learning rate *α* determines the weighting of new experience relative to existing knowledge. In deterministic physical systems, we set *α* = 1 to allow rapid convergence toward optimal policies. The discount factor *γ* reflects the swimmer’s emphasis on long-term displacement; small *γ* encourages exploiting immediate gains, while large *γ* promotes strategic, multi-step propulsion cycles.

The swimmer’s position is evaluated based on its body centroid, calculated as the mean position of all spheres:

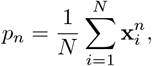

where 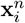 is the position of sphere *i* at learning step *n*. The displacement after one action is:

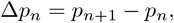

and the immediate reward is defined as:

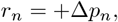

encouraging net positive displacement in the swimmer’s preferred direction.

To balance exploration and exploitation during training, we employ the epsilon-greedy strategy. At each step, the swimmer selects a random action with probability *ϵ*, encouraging exploration of new state-action pairs. With probability 1 − *ϵ*, it exploits its current knowledge by choosing the action maximizing *Q*(*s, a*). For more complex systems the parameter *ϵ* typically sets to a high number initially with decaying over time, promoting thorough high random exploration initially and convergence toward optimal policies as learning progresses.

This RL framework enables the swimmer to autonomously discover optimal sequences of deformations that leverage hydrodynamic interactions to achieve efficient locomotion in viscous, low Reynolds number environments.

## 3 Results and Discussion

In this section, we present the results of reinforcement learning (RL) applied to a variety of microswimmer models, clearly distinguishing between reproduced benchmarks and novel contributions. We begin by revisiting the classical three-sphere swimmer, originally proposed and previously studied under RL frameworks, to validate the correctness and reliability of our implementation.

Next, we consider the Double Spring swimmer, which replaces rigid linkers with oscillatory springs. Although this physical model has been introduced in earlier works, RL has not yet been applied to it. We extend it by incorporating phase-controlled oscillatory deformations and analyze the emergence of efficient propulsion strategies. The Charged Spring swimmer represents a novel contribution in this study, where electrostatic interactions between charged spheres are introduced to mimic chemotactic behavior in biological systems. We then generalize the linear architecture to *N*-sphere swimmers. While such models have been explored in prior literature, including under RL, we reproduce key results as a validation of the scalability of our framework.

Following the linear systems, we explore geometrically richer designs. The Triangular swimmer, although theoretically and numerically studied before, is investigated here for the first time using reinforcement learning. The Plus-like and Cage-like swimmers, proposed and developed in this work, represent novel 2D and 3D architectures, respectively. These models demonstrate how geometric complexity and actuation constraints influence learning dynamics and propulsion efficiency.

### 3.1 Three-sphere swimmer and its extensions

The classical three-sphere swimmer, first introduced as a minimal model for low Reynolds number propulsion [19], has been extensively analyzed in theoretical studies, including its hydrodynamic behavior and symmetry-breaking requirements [13]. More recently, reinforcement learning approaches have also been applied to this model to explore its capacity for data-driven propulsion discovery [22].

To validate our RL implementation and ensure consistency with prior results, we first trained the classical three-sphere swimmer using a constant exploration rate *ϵ* = 0.1, learning rate *α* = 1, and discount factor Γ = 0.6. The agent successfully discovered an efficient propulsion strategy within approximately 22 learning cycles, each consisting of 100 steps. Analysis of the learned state transition sequence revealed that the swimmer reproduced the well-established optimal motion cycle proposed in earlier studies. This alignment confirms both the validity of the reinforcement learning framework and the physical consistency of the model. Figure 3a shows the evolution of displacement over learning cycles, and Figure 3b illustrates the optimal cycle structure discovered by the agent.

**Figure 1.**
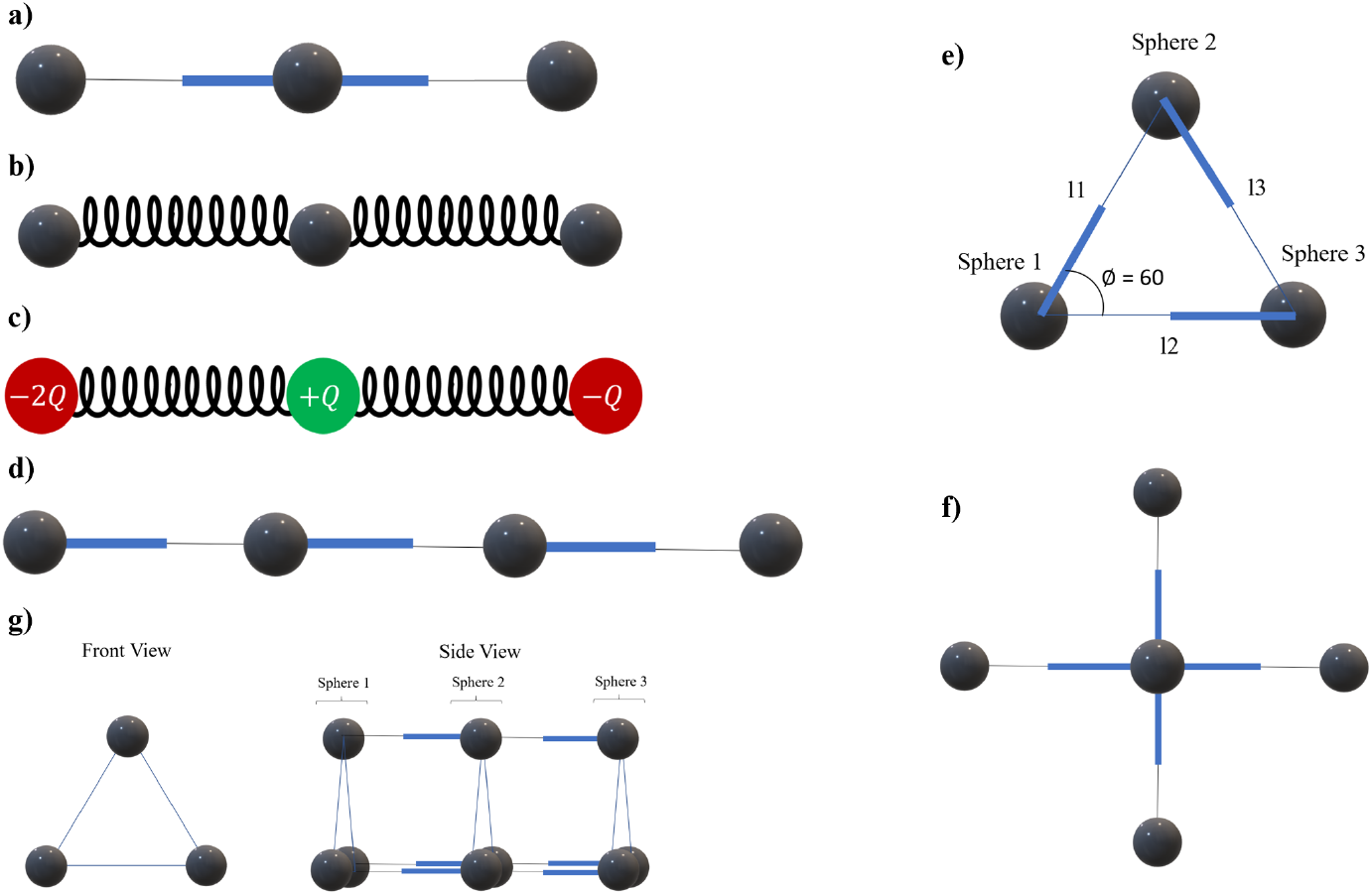
Illustration of swimmer architectures studied in this work: (a) Classical linear three-sphere swimmer; (b) Double spring model with oscillatory linkers; (c) Charged spring model with electrostatic interactions; (d) Extended linear 4-sphere swimmer; (e) Triangular swimmer enabling planar and rotational motion; (f) Plus-like swimmer with orthogonal linker arrangement; (g) Cage-like swimmer with triangular cross-section and constrained linear locomotion.

**Figure 2.**
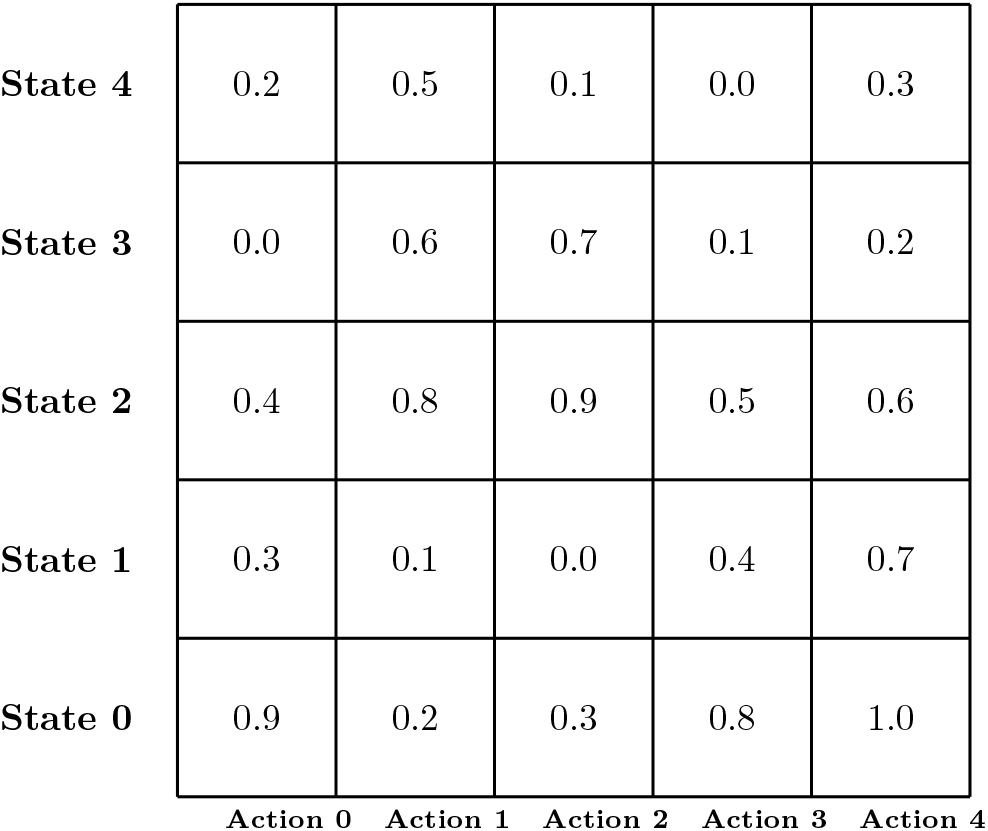
Example of visualization of the Q-matrix after training, representing learned expected rewards for each state-action pair. Higher values indicate actions contributing to efficient propulsion strategies.

**Figure 3.**
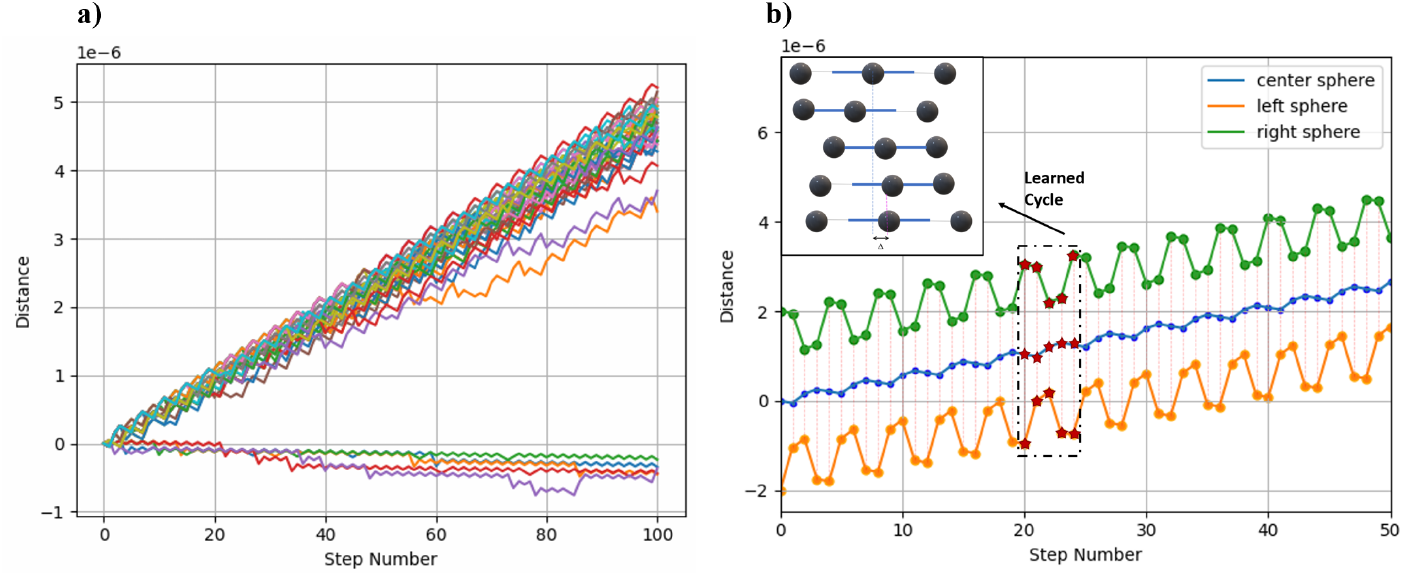
a) Total displacement versus training steps. b) Learned optimal motion cycle for the three-sphere swimmer, matching the theoretical swimming sequence proposed in previous studies.

While the classical three-sphere swimmer provides a minimal framework for studying propulsion at low Reynolds numbers, it differs fundamentally from many biological microswimmers, such as cyanobacteria, due to its use of rigid, discrete linker deformations. Theoretical and numerical analyses have previously explored generalized versions of this swimmer incorporating continuous or elastic deformations [13, 29, 30], but to the best of our knowledge, reinforcement learning has not yet been applied to such physically enriched models to evaluate their autonomous learning capabilities.

To bridge this gap and better approximate realistic, continuous deformation patterns observed in nature, we modified the swimmer by replacing the rigid linkers with elastic springs of identical spring constants *k*_1_ = *k*_2_ = 10^−4^. These springs oscillate around their equilibrium lengths at a fixed frequency of *ω* = 1 Hz, mimicking sinusoidal deformations. This setup allows us to investigate how a microswimmer can learn to exploit phase differences between the oscillations to achieve propulsion, similar to traveling wave dynamics seen in biological systems.

This modification, however, raises an important conceptual question regarding the interpretation of actions within the reinforcement learning framework. To address this, we fixed the phase of one spring to *ϕ*_1_ = 0 and allowed the second spring to oscillate with a tunable phase difference *ϕ*_2_. To discretize the action space while maintaining manageable computational complexity, the phase difference *ϕ*_2_ was divided uniformly into 10 possible values within the range [0, *π*]. Consequently, each action corresponds to a controlled transition of the system’s phase configuration from one discrete phase difference to another. For reward calculation, we implemented a cyclic deformation approach: after each action selection, the swimmer undergoes a complete oscillation cycle, and the resulting net displacement serves as the reward signal for the agent.

Given the symmetry of the swimmer, one can anticipate that optimal propulsion should emerge when the phase difference approximates Δ*ϕ* = *π/*2, resembling the traveling longitudinal wave propagation observed in biological systems such as nematode locomotion, cyanobacterial gliding [9], and amoeboid crawling [2].

Due to the increased complexity of the system’s continuous deformation space, we adopted a decaying exploration strategy with an initial exploration rate of *ϵ* = 0.9, final exploration rate *ϵ*_final_ = 0.2, and decay rate *λ* = 0.99. The learning rate was set to *α* = 0.9, and the discount factor to Γ = 0.6. These parameters provided a balance between sufficient exploration of the complex state space and efficient convergence. Our simulations confirmed the biological analogy. As shown in Figure 4, over 200 training cycles of 100 steps each, the swimmer gradually optimized its motion, converging to an average phase difference of approximately 0.72*π*. This slight deviation from the theoretical optimum likely results from simplified numerical approximations in our current implementation.

**Figure 4.**
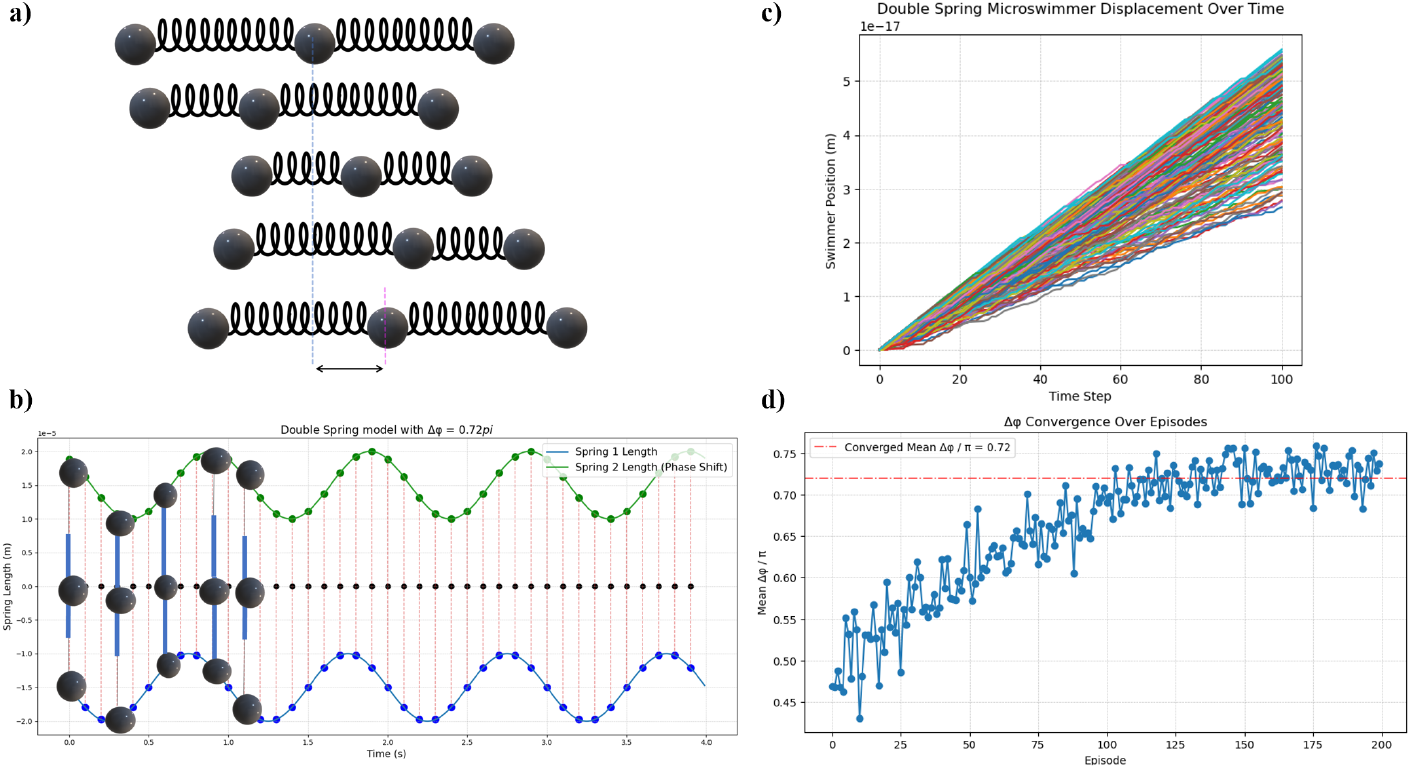
(a) Learned optimal propulsion cycle based on controlled phase differences; (b) Oscillatory deformation of the springs over time, showing convergence to a periodic pattern approximating the classic longitudinal traveling wave; (c) Net displacement of the swimmer versus time across all learning cycles, indicating improvement in propulsion efficiency; (d) Convergence of the phase difference Δ*ϕ* between the springs toward the optimal value near *π/*2.

To further approximate realistic microswimmers exhibiting chemotaxis or bio-inspired behavior, we extended the oscillatory three-sphere swimmer by introducing electrostatic interactions. Specifically, the left sphere was assigned a charge of −2*Q*, the central sphere +*Q*, and the right sphere −*Q*, with *Q* = 10^−14^ C. The equilibrium lengths of the springs and the positions of the spheres were carefully tuned such that, at the system’s rest configuration, the sum of all elastic and electrostatic forces vanishes, ensuring a stable global equilibrium.

This configuration introduces an interesting asymmetry: during the first step of each oscillatory cycle, the positively charged central sphere experiences an enhanced attractive force toward the left sphere, whose negative charge is twice that of the right sphere. As a result, the deformation of the springs during this phase occurs more rapidly, leading to increased net displacement per action in comparison to the uncharged model.

We applied the same phase-based reinforcement learning strategy as before, discretizing the phase difference Δ*ϕ* into 10 values within [0, *π*]. We again employed a decaying exploration rate starting from *ϵ* = 0.9, decaying exponentially to *ϵ*_final_ = 0.2 with a decay factor of 0.99, alongside learning rate *α* = 0.9 and discount factor Γ = 0.6.

Remarkably, as shown in Figure 5, the swimmer successfully converged to an optimal phase difference of approximately 0.5145*π* within just 30 training cycles of 30 steps each. This value lies very close to the theoretical optimum of Δ*ϕ* = *π/*2, consistent with prior theoretical models and observations of biological microswimmers. We hypothesize that the improved convergence and strong agreement with biological systems stem from the introduction of electrostatic interactions, which bring the model closer to real-world physical conditions.

**Figure 5.**
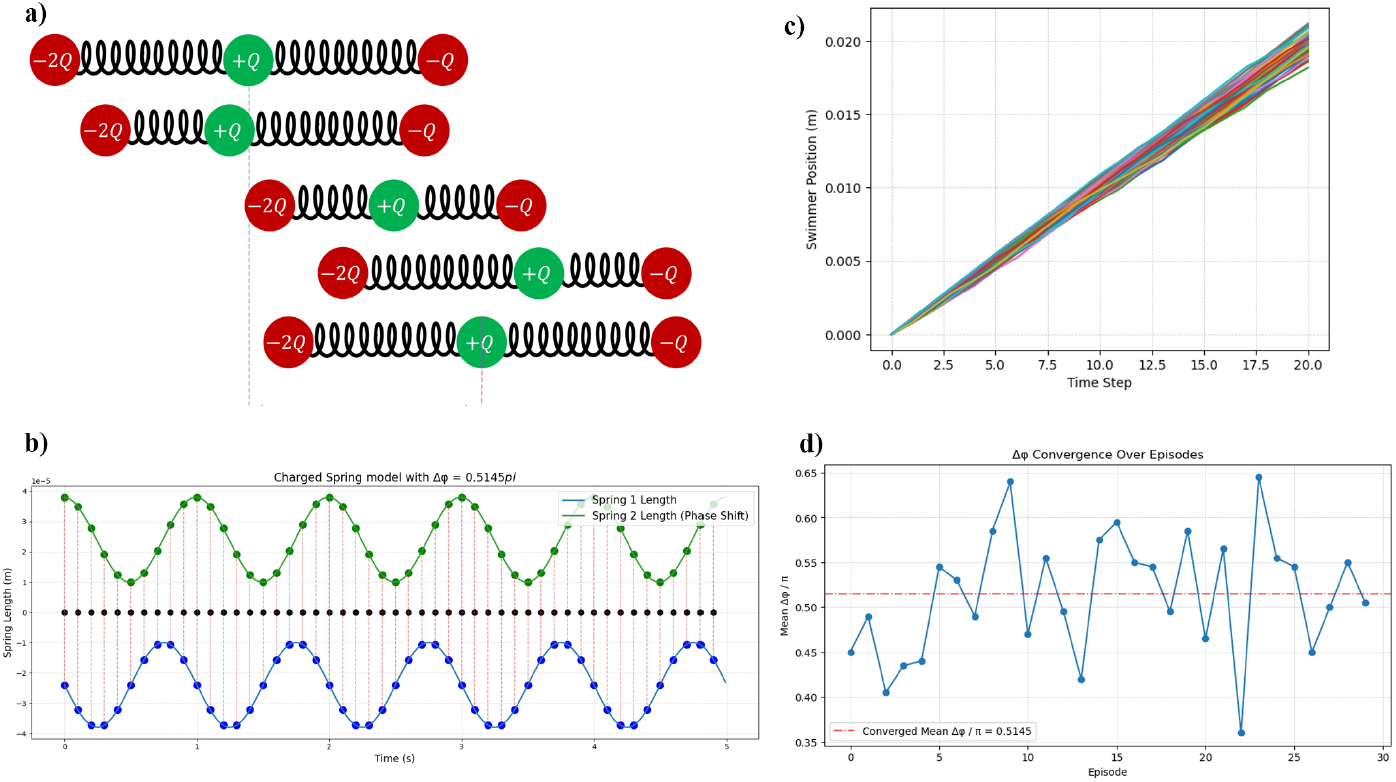
(a) Learned optimal propulsion cycle (b) Oscillatory deformation of the springs over time, showing a better and faster convergence to a periodic pattern approximating the classic longitudinal traveling wave; (c) Net displacement of the swimmer versus time across all learning cycles, indicating improvement in propulsion efficiency; (d) Convergence of the phase difference Δ*ϕ* between the springs toward the optimal value near *π/*2.

It is noteworthy that, as observed in our simulations, the displacement per learning cycle of the Double spring swimmer is significantly lower than that of the classical three-sphere swimmer, despite the biological realism introduced by continuous deformations. In contrast, the Charged spring swimmer demonstrates a considerable increase in propulsion efficiency, achieving greater net displacement compared to both the classical and Double spring models within fewer learning cycles. We attribute this enhanced performance to the presence of electrostatic interactions, which introduce additional, naturally occurring forces that assist deformation and propulsion processes. This finding is consistent with the strategies employed by biological microsystems, where cells and microorganisms exploit electrostatic potential differences, membrane polarization, or ionic gradients to regulate movement, deformation, and transport. The improved efficiency observed in our charged swimmer model supports the hypothesis that incorporating electrostatic interactions brings artificial microswimmers closer to mimicking nature’s highly optimized locomotion mechanisms.

In other words, the improved efficiency observed in our charged swimmer model highlights the evolutionary intelligence of natural systems, which have optimized their locomotion and transport mechanisms over millions of years by leveraging fundamental physical principles such as electrostatics to overcome constraints at microscopic scales.

### 3.2 Generalized *N*-sphere swimmer

The linear *N*-sphere swimmer, as a direct generalization of the classical three-sphere model, has been extensively studied to investigate scalable propulsion strategies at low Reynolds numbers [1, 19]. Prior research has explored both analytical and numerical aspects of these extended models, and reinforcement learning has also been applied in select studies to investigate the emergence of optimal swimming policies in such systems [15].

To validate the consistency and correctness of our reinforcement learning implementation, we reproduced this model as a benchmark case. Specifically, we considered a swimmer composed of *n* = 4 spheres connected in a linear configuration. The agent was trained using 80 learning episodes, each consisting of 150 steps, with hyperparameters set to *α* = 1, Γ = 0.8, and a constant exploration rate of *ϵ* = 0.05. This benchmark serves both as a validation of our framework and a baseline for comparison with more complex swimmer architectures.

The results are summarized in Figure 6, demonstrating successful convergence of the propulsion strategy and displacement improvements in agreement with prior studies. These findings confirm the validity of our code and provide a robust baseline for exploring more complex swimmer architectures.

**Figure 6.**
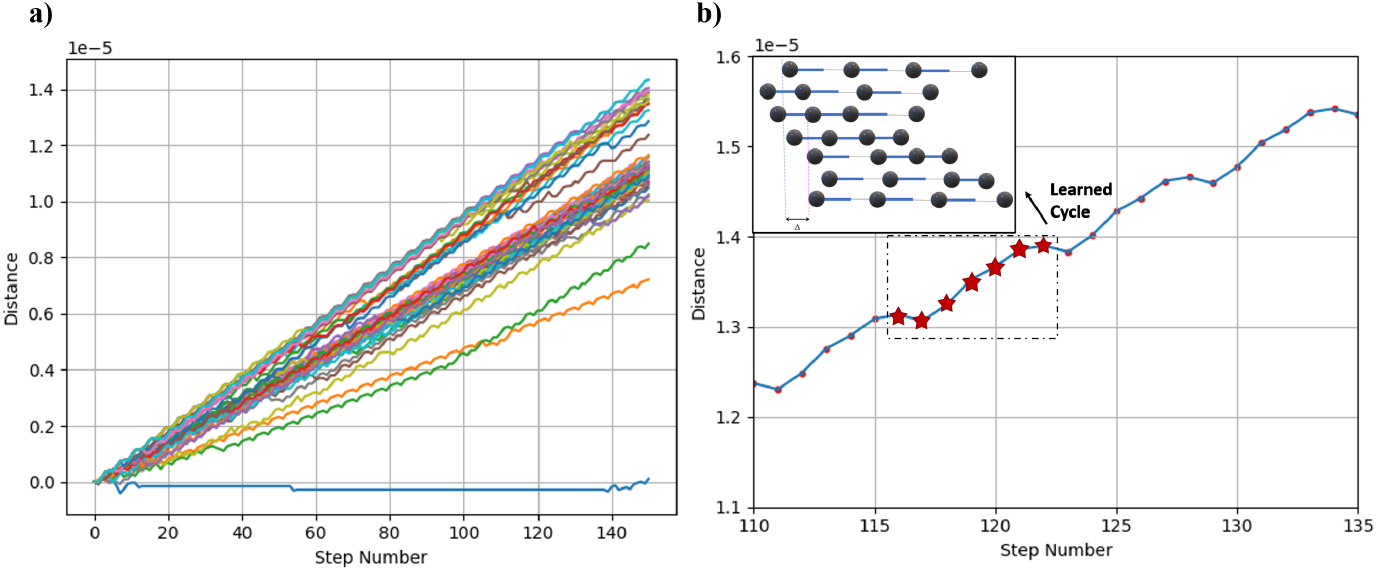
a) Total displacement versus training steps. b) Learned optimal motion cycle for the 4-sphere swimmer, matching the theoretical swimming sequence proposed in previous studies.

As shown in our simulations, increasing the number of spheres in the swimmer generally leads to enhanced propulsion performance, with higher net displacement per cycle. However, this improvement comes at the cost of increased action-state space complexity, which requires more learning steps to converge toward an optimal swimming strategy.

Additionally, it is important to consider that as the number of spheres increases, the overall size of the microswimmer grows. From an experimental standpoint, constructing such swimmers with a larger number of spheres would require using smaller components to ensure the system remains within the low Reynolds number regime, where viscous forces dominate. As expected, the learned optimal propulsion patterns for the *N*-sphere swimmer naturally generalize the longitudinal traveling wave strategy previously proposed for the three-sphere model, confirming both the scalability of the mechanism and the effectiveness of our learning framework.

In the following section, we shift our focus to the triangular swimmer, which, due to its unique geometric configuration, introduces distinct dynamical behaviors compared to the linear models discussed so far.

### 3.3 Triangular swimmer

The triangular microswimmer represents one of the most established theoretical models for exploring locomotion at low Reynolds numbers, especially for understanding planar, two-dimensional swimming strategies [8, 24]. While its hydrodynamics and deterministic swimming patterns have been extensively analyzed, to the best of our knowledge, this is the first time reinforcement learning (RL) has been applied to autonomously optimize its propulsion.

The key distinction of the triangular swimmer lies in its geometry: composed of three spheres connected via deformable linkers forming a closed triangular structure, this swimmer can generate net displacement in two dimensions by cyclically contracting and extending its linkers. Additionally, due to asymmetries in deformation, it exhibits small rotational displacements about its center of mass, typically on the order of 10^−19^ radians per cycle under standard parameter regimes.

To better visualize the rotational dynamics during simulations, we amplified the recorded angular displacements by a factor of 10^9^ for clarity in the plots.

The RL training was performed over 30 learning cycles, each consisting of 50 steps. The hyperparameters were set to *α* = 0.8, Γ = 1, and a constant exploration rate *ϵ* = 0.3.

The following results, summarized in Figure 7, demonstrate the system’s ability to autonomously discover efficient swimming patterns in two-dimensional space, along with small but consistent rotational drift, confirming the model’s enhanced locomotion flexibility compared to linear swimmers.

**Figure 7.**
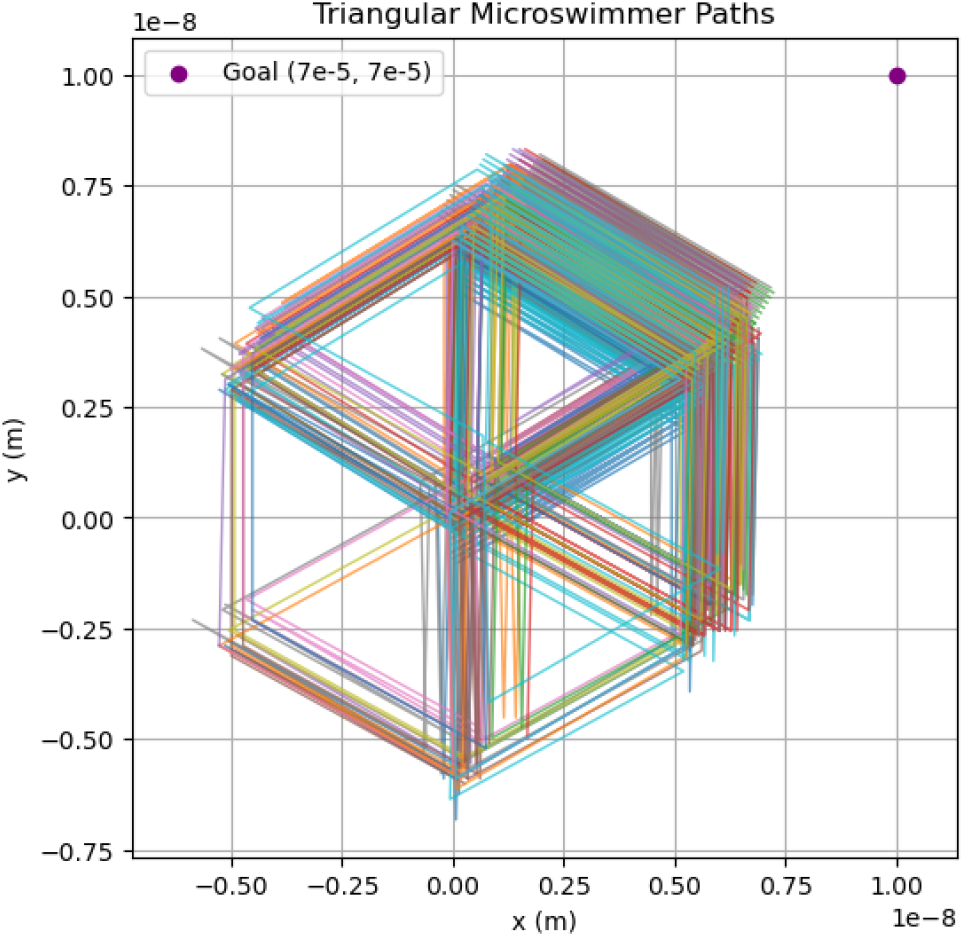
Overall trajectory of the triangular microswimmer during the 30 learning propulsion cycles.

As observed in the simulations, the triangular swimmer moves toward its target following an overall hexagonal trajectory with relatively low propulsion efficiency. Nevertheless, its small rotational drift enables it to escape purely reciprocal cycles and achieve net displacement. We hypothesize that, similar to the mechanisms discussed in [8], incorporating stochastic effects such as controlled diffusion into the system may allow the swimmer to explore alternative, potentially more efficient locomotion strategies beyond the current deterministic framework.

Although we are not entirely certain about this result due to the complexity of solving the governing equations, which compelled us to apply approximations, this outcome is obtained based on a first-order approximation of the hydrodynamic equations and should be interpreted with caution.

### 3.4 Plus-like swimmer

The next swimmer architecture introduced in this study is an entirely novel design conceptualized and developed by our group. Inspired by the simplicity and efficiency of the Najafi-Golestanian three-sphere swimmer [20], we extended the concept into a more complex structure resembling two orthogonal three-sphere swimmers superimposed with a 90^°^ angular offset, forming what we term the “Plus-like” swimmer.

This swimmer consists of five spheres connected via four independent linkers aligned along the horizontal and vertical axes. To maintain manageable action-state complexity, the swimmer is constrained such that only one linker may deform per action. Consequently, it possesses the capability to generate controlled displacements along both Cartesian axes, offering true two-dimensional locomotion. To facilitate target-oriented navigation, we modified the reward function compared to previous models. Instead of assigning displacement as a reward, we defined the reward as the negative of the swimmer’s Euclidean distance to the target after each action. In this formulation, minimizing distance translates to maximizing reward, effectively encouraging the swimmer to continuously reduce its distance to the goal.

We hypothesized that, under this setup, when the target lies directly along the horizontal or vertical axis, the swimmer would adopt propulsion strategies analogous to the classical three-sphere swimmer. Conversely, for diagonal targets, we expected the swimmer to alternate its axis-specific deformations to approach the goal along a diagonal trajectory.

To test this, we trained the swimmer over 1500 learning cycles, each consisting of 2000 steps, using hyperparameters *α* = 1, Γ = 0.8, and a constant exploration rate *ϵ* = 0.15.

Following training, we validated the learned policy by deploying the updated *Q*-matrix in scenarios with purely horizontal and vertical targets. The swimmer successfully exhibited behavior consistent with our predictions, achieving efficient, goal-directed locomotion along both axes, as illustrated in Figure 8.

**Figure 8.**
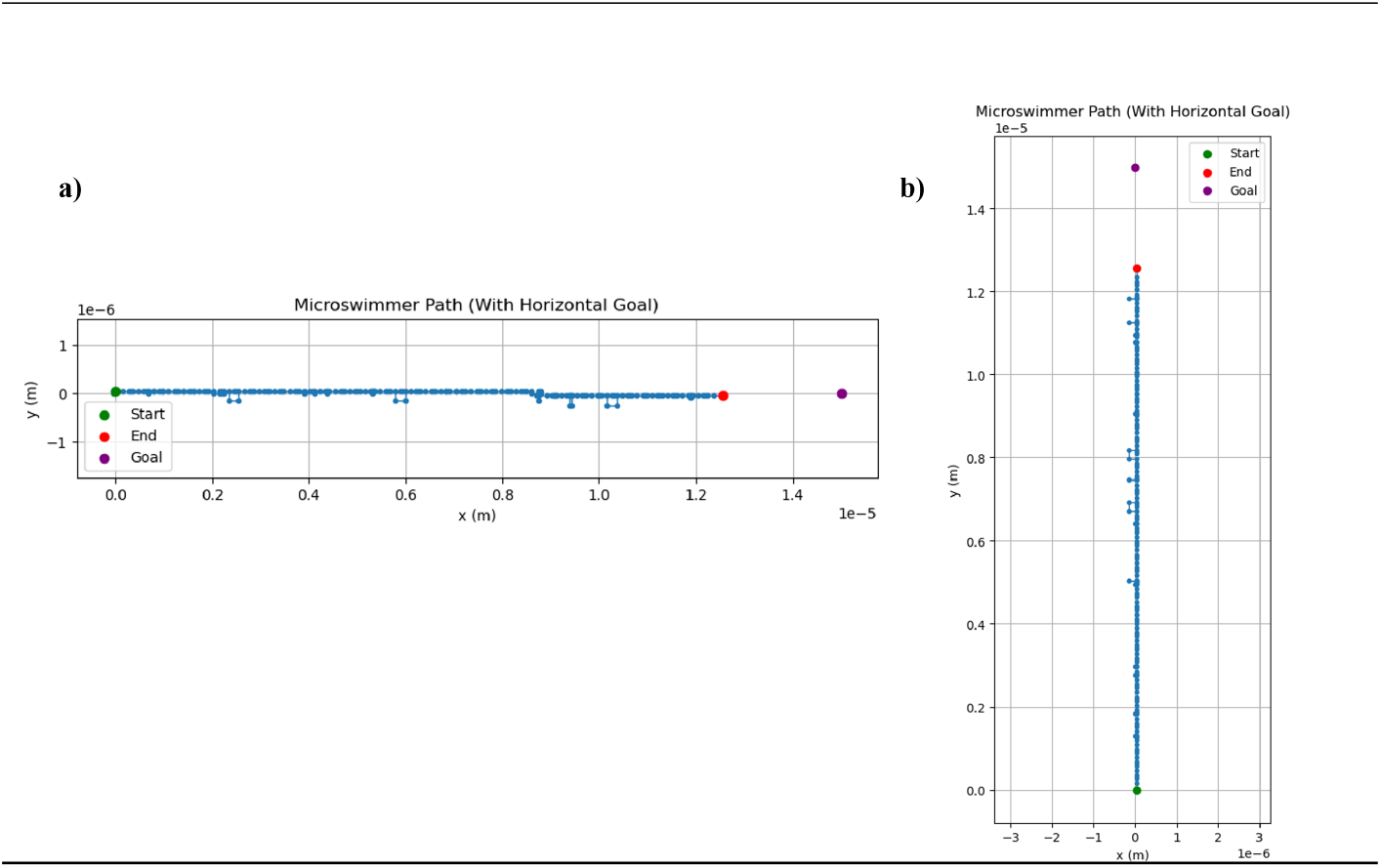
Overall trajectory of the Plus-like microswimmer with both horizontal and vertical goals after the full learning.

Subsequently, we retrained the Plus-like swimmer to target a diagonal goal using the same configuration of 1500 learning cycles, each with 2000 steps. The complete trajectory of the swimmer under this setup is presented in Figure 9, demonstrating its ability to coordinate deformations along both axes to achieve efficient diagonal locomotion toward the target.

**Figure 9.**
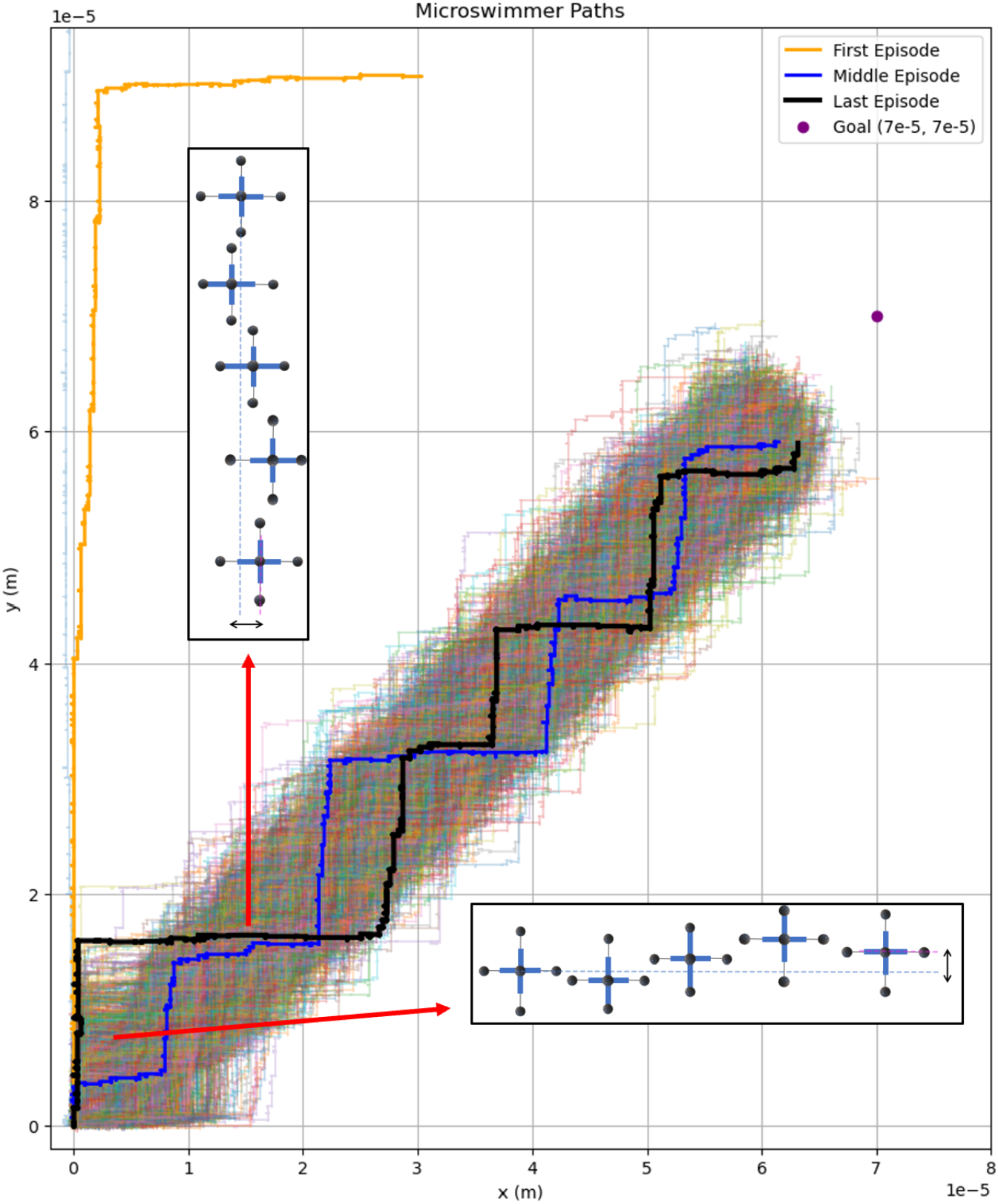
Overall trajectories of the Plus-like microswimmer over 1500 cycles with a diagonal goal.

As illustrated in Figure 9, the swimmer initially attempts to reach the diagonal target by separately executing purely vertical and purely horizontal movements. However, due to the limited number of steps and the inefficiency of strictly axis-aligned strategies for diagonal targets, the swimmer fails to approach the goal effectively during the early stages of learning. Subsequently, the swimmer explores stepwise trajectories resembling staircase-like paths, composed of alternating horizontal and vertical segments. This hybrid strategy enables gradual progress toward the diagonal goal.

It is noteworthy that during the initial learning cycles, these stepwise trajectories exhibit significant irregularities and noise, with inconsistent segment lengths and misaligned angular transitions. However, as training progresses, the swimmer refines its policy, producing increasingly orthogonal and smoother staircase patterns with longer, more efficient steps along both axes. These observations highlight the capability of the Plus-like swimmer to autonomously discover and optimize complex locomotion strategies suitable for arbitrary two-dimensional targets. Moreover, the gradual improvement in trajectory regularity reflects the effectiveness of reinforcement learning in refining control policies, even within high-dimensional action spaces.

Ultimately, to progress toward practical applications such as targeted drug delivery, we introduce a novel three-dimensional swimmer architecture, termed the Cage-like swimmer.

### 3.5 Three-dimensional Cage-like swimmer

Inspired by biological microscale transport mechanisms, the Cage-like swimmer introduced in this work is a novel three-dimensional microswimmer designed to encapsulate and transport cargo while navigating viscous environments. Its structure consists of a triangular cross-sectional frame formed by three spheres connected via deformable linkers, resembling the planar triangular swimmer, but further augmented with additional linkers to form a closed 3D cage-like geometry. This enclosure not only enables the swimmer to mimic encapsulation behaviors observed in natural systems but also introduces enhanced spatial control, potentially allowing movement along all three spatial axes.

Although similar concepts have appeared in prior studies, such as microswimmers navigating within droplets [23], enclosed microcages [5], and 3D-printed biodegradable swimmers [4], those implementations typically involve complex materials or physical mechanisms. In contrast, the model proposed here is deliberately minimal in both structure and actuation rules, making it ideal for computational modeling and reinforcement learning experiments. Crucially, the primary aim of this investigation is not simply to design a swimmer with optimal performance, but to understand *“how a minimal and biologically inspired system can learn to swim”* through interaction with its environment. By embedding the swimmer in a reward-driven reinforcement learning (RL) framework, we enable it to autonomously discover propulsion strategies adapted to its geometry and constraints. We also imposed specific simplifications during training. First, the swimmer’s motion was constrained to one-dimensional translation; the triangular cross-section was not utilized for rotational or lateral movement. Second, we employed a symmetric actuation scheme: the three front-facing linkers deform synchronously, and likewise the three rear linkers, alternating over time. This simplification reduced the action space while preserving enough degrees of freedom for meaningful motion.

Despite these constraints, the swimmer successfully learned an efficient propulsion policy. Given its resemblance to the classical three-sphere swimmer, we hypothesized that it would converge toward a generalized traveling-wave-like deformation strategy, with approximately triple the displacement per cycle. Indeed, training the model for 50 episodes of 100 steps each, with parameters *α* = 1, Γ = 0.6, and *ϵ* = 0.1, resulted in a convergent Q-matrix and consistent displacement trajectories. These findings confirm that even under strong structural and control limitations, the Cage-like swimmer can learn to propel effectively through viscous media (see Figure 10).

**Figure 10.**
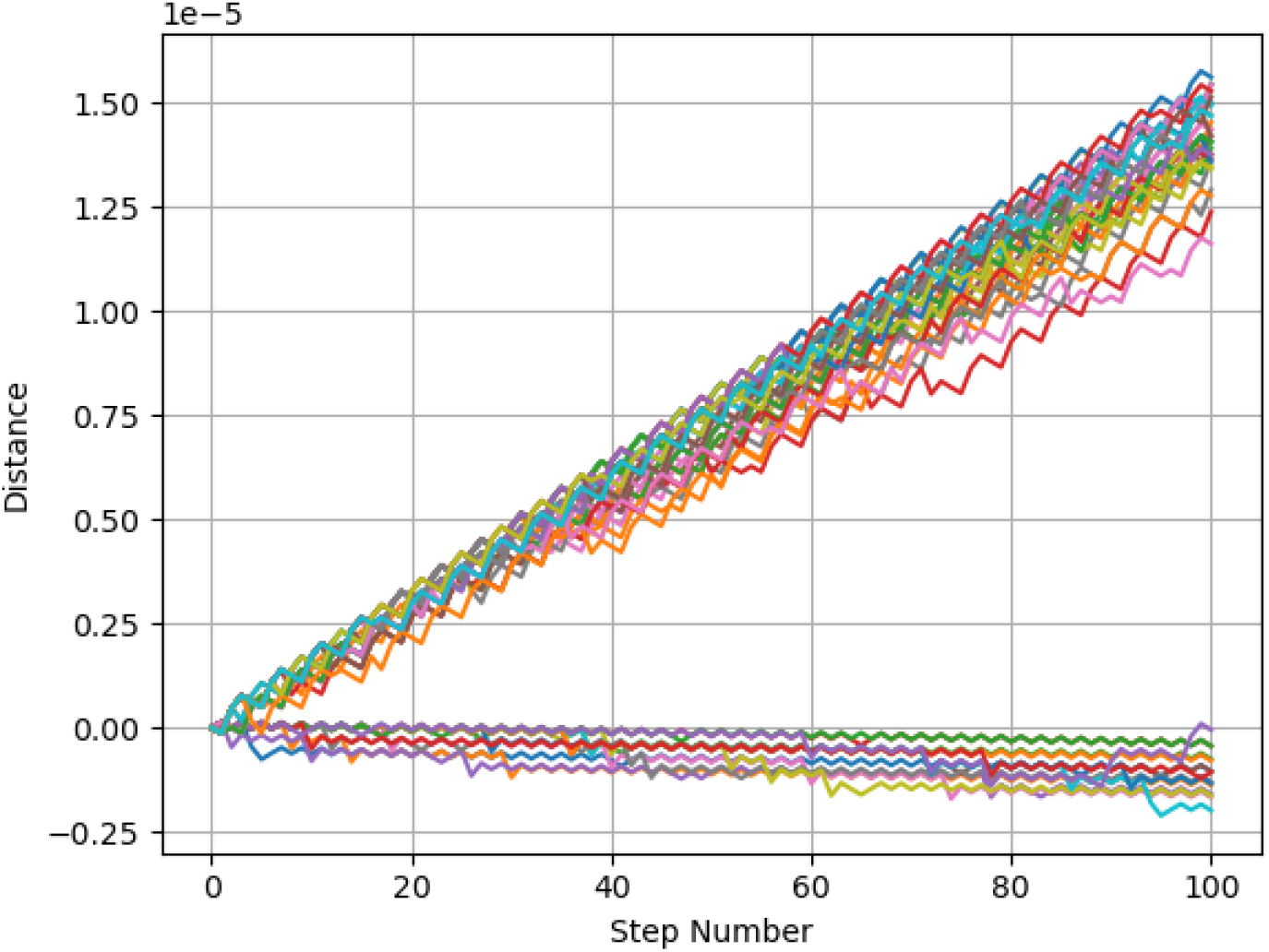
Total displacement versus training steps.

As hypothesized, and as clearly observed in the displacement trajectory (Figure 10), the net distance traveled per optimal learned cycle is approximately three times that of the classical three-sphere swimmer. This outcome highlights the significant impact of the swimmer’s geometric augmentation, despite the actuation constraints imposed during training.

Having explored the reinforcement learning behavior across a wide variety of swimmer geometries, we now provide a comparative analysis to distill key insights from the experiments. We analyze swimmer performance, learning efficiency, and locomotion strategies, identifying the trade-offs between structural complexity and learning convergence. Furthermore, we discuss the broader implications of these findings for designing efficient artificial microswimmers and outline directions for future research.

## 4. Discussion and Conclusion

In this study, we systematically explored how diverse microswimmer architectures can autonomously learn efficient propulsion strategies in low Reynolds number environments via reinforcement learning. Our results span both classical and novel swimmer models, enabling comparative insights across levels of geometric and physical complexity.

### 4.1 Comparative analysis of swimmer performance

As observed in our simulations, the swimming performance of linear microswimmers improves with an increasing number of spheres. This is consistent with previous theoretical predictions [15, 19, 22], where the enhanced spatial degrees of freedom allow more effective hydrodynamic coordination. However, increasing the number of spheres also increases the physical size of the swimmer. Since the Reynolds number depends on both velocity and length scale, it is essential to verify whether such extended swimmers still operate within the Stokes flow regime.

In terms of learning dynamics, simpler models with fewer spheres (e.g., the classical three-sphere swimmer) converged more rapidly to effective policies, even with a constant exploration rate *ϵ*, indicating a lower complexity in their action-state space. For these models, we avoided decaying *ϵ* to reduce training overhead, yet convergence was swift and consistent.

Replacing rigid linkers with elastic springs marked a shift toward more biologically realistic deformations. This oscillatory actuation strategy introduced phase-dependent displacement patterns, and as expected from prior theoretical analyses [2, 9, 13], we found that optimal propulsion tends to emerge near a phase difference of Δ*ϕ* ≈ *π/*2. This suggests that our learning framework not only captures efficient behavior but converges to physically interpretable, biologically-inspired dynamics.

Furthermore, the introduction of electrostatic interactions through charged spheres brought the model even closer to realistic chemotactic microrobots. While the learned swimming behavior still resembled traveling-wave-like deformations, the total displacement per cycle increased. Although this result is currently speculative, we hypothesize that the additional electrostatic forces assist propulsion, effectively mimicking biological strategies where internal electric potentials are known to mediate cellular motion [11]. These advanced models required longer training cycles to converge due to their increased complexity and continuous interaction potentials.

### 4.2 Insights from non-linear swimmer geometries

Our investigation into non-linear swimmer geometries, such as the triangular, plus-like, and cage-like models, further demonstrates the flexibility of reinforcement learning in discovering viable propulsion strategies beyond simple linear chains.

The triangular swimmer, while limited in efficiency, leveraged its small rotational degrees of freedom to explore a wider space of configurations. The learned motion resembled a hexagonal path with slight angular drift per cycle. This residual rotational motion enabled incremental displacement over time, although the swimming speed remained modest. Previous studies have suggested that adding stochastic rotational diffusion to such swimmers may improve their search capabilities or directional exploration [8], a direction we did not pursue here but consider promising for future work.

The plus-like swimmer, a novel architecture introduced in this work, required significantly longer training to converge. Owing to its orthogonal design and symmetric actuation constraints, the learned locomotion strategy emerged as a staircase-like motion toward its target. This pattern consisted of alternating vertical and horizontal displacements. Early training episodes exhibited irregular trajectories, but over time the swimmer learned to produce smoother, longer steps, demonstrating both local exploration and policy refinement.

In the cage-like swimmer, another novel contribution of this paper, we constrained the three-dimensional geometry to one-dimensional linear motion by enforcing symmetric linker actuation. Interestingly, the learned strategy closely mirrored that of the classical three-sphere swimmer. However, due to its extended structure, effectively consisting of three parallel three-sphere units, the net displacement per cycle was approximately three times greater. This highlights the cumulative power of modular swimmer design and suggests that complex swimmers can leverage repeated substructures to amplify motion.

### 4.3 Limitations and Future Directions

While our framework successfully demonstrates learning across a wide range of swimmer designs, several limitations merit discussion. First, we employed a relatively simple Q-learning algorithm in a discretized configuration space. More advanced reinforcement learning methods, such as Deep Q-Networks (DQN), actor-critic algorithms, or policy gradient methods, could enable learning in continuous, high-dimensional control spaces and improve scalability and generalization [18].

Second, our simulations did not incorporate environmental stochasticity or physical perturbations. Realistic microswimmer behavior is strongly influenced by thermal fluctuations (Brownian motion), rotational diffusion, and hydrodynamic noise, particularly at micrometer scales. Including thermal noise and diffusion in future models would allow us to better assess the robustness of learned policies and understand how swimmers navigate under uncertainty [8, 13, 15].

Third, we assumed idealized, unbounded fluid domains without boundaries or obstacles. However, interactions with confining geometries, channel walls, or crowded environments can dramatically alter swimming efficiency and directional persistence [31]. Extending our models to account for such constraints could lead to new insights relevant for applications in microfluidic channels, biological tissues, or synthetic scaffolds.

In terms of system complexity, our current work focused on single-agent learning. Multi-agent extensions may reveal emergent collective behaviors such as swarming, alignment, or cooperative transport, as observed in both bacterial colonies and synthetic microrobot swarms [17].

Finally, we constrained the locomotion of geometrically complex swimmers, such as the cage-like swimmer, to simplified 1D motion to facilitate tractable learning. Relaxing these constraints in future studies may uncover richer behavioral repertoires, including rotation, steering, and full 3D trajectory planning.

Future work will thus explore:

- Employing deep reinforcement learning for continuous control and high-dimensional policy spaces.
- Adding environmental noise, such as thermal fluctuations and Brownian motion.
- Modeling interactions with physical boundaries and obstacles.
- Investigating cooperative learning among multiple swimmers.
- Allowing unconstrained 3D motion and dynamic shape adaptation.

### 4.4 Concluding remarks

Our results demonstrate that minimal microswimmer models, when equipped with basic reinforcement learning capabilities, can autonomously discover propulsion strategies that align with biological behaviors and theoretical predictions. By progressing from classical to biologically inspired and novel geometries, we show that learning-based frameworks are a powerful tool for probing microscale locomotion. The simplicity of our swimmer designs and learning mechanisms also make this approach well-suited for computational exploration and could inform future experimental realizations of smart, adaptive microrobots in viscous environments.

## Acknowledgements

I would like to express my deepest gratitude to Prof. S.Mehdi Vaez Alaei for his invaluable guidance, mentorship, and continued support throughout the course of this research. His insightful feedback, rigorous scientific approach, and encouraging supervision were instrumental in shaping the direction and quality of this work. I am especially grateful for the intellectual freedom he granted me, which allowed for the creative development of the ideas presented in this study.

## Data and Code Availability

For code and implementation details related to this work, interested readers are encouraged to contact the author at.

